# Dynamic coastal pelagic habitat drives rapid changes in growth and condition of juvenile sockeye salmon (*Oncorhynchus nerka*) during early marine migration

**DOI:** 10.1101/2022.03.21.484660

**Authors:** Jessica Garzke, Ian Forster, Sean Godwin, Brett T. Johnson, Martin Krkosek, Natalie Mahara, Evgeny A. Pakhomov, Luke A. Rogers, Brian P.V. Hunt

## Abstract

Migrating marine taxa encounter diverse habitats that differ environmentally and in foraging conditions over a range of spatial scales. We examined body (RNA/DNA, length-weight residuals) and nutritional (fatty acid composition) condition of juvenile sockeye salmon (*Oncorhynchus nerka*) in British Columbia, while migrating through varied oceanographically waters. Fish were sampled in the stratified northern Strait of Georgia (NSoG); the highly mixed Johnstone Strait (JS); and the transitional zone of Queen Charlotte Strait (QCS). In 2015, body and nutritional condition were high in the NSoG and responded rapidly to reach lowest levels in JS with its low prey availability, and showing signs of compensatory growth in QCS. In 2016, juvenile salmon had significantly lower condition in the NSoG than in 2015, although zooplankton biomass was similar, condition remained low in JS, and no compensatory growth was observed in QCS. We provide evidence that differences in juvenile salmon condition between the two years being due to changes in the food quality available to juvenile fish. Further, we propose that the TGH needs to be extended to incorporate food quality as a parameter to understand changes in fish condition and survival between years.

## Introduction

Pacific salmon (*Oncorhynchus* spp.) are a group of anadromous species that experience extreme spatial variations in ecosystem attributes during their early marine life, during which they typically migrate from estuaries to the coastal ocean and then to the open ocean. Pacific salmon can experience high mortality during their first months at sea, with mortalities of up to 80-90% estimated for tagged salmon (Welch et al., 2011). It has been hypothesized that growth and mortality rates during this phase may contribute to interannual variability and long-term population trends of salmon stocks (Friedland et al., 2000; Mueter et al., 2005). Although there are likely numerous factors that influence juvenile salmon survival during their early marine phase, correlative evidence suggests that feeding conditions play an important role (Beamish et al., 2012; Duffy and Beauchamp, 2011). Through impacts on growth, feeding conditions have the potential to affect vulnerability to predation (Gliwicz, 2002), ability to survive stressful ocean conditions (Lasker, 1978), and the body condition required to survive their first ocean winter (Beamish and Mahnken, 2001).

Recent research from North America’s west coast has drawn attention to the potential influence of small-scale coastal marine conditions on juvenile salmon survival (Ferriss et al., 2014; Hunt et al., 2018; Journey et al., 2018; McKinnell et al., 2014). Outmigrating juvenile Fraser River sockeye (*Oncorhynchus nerka)* first enter the ocean into the stratified, comparatively warm, and highly productive Strait of Georgia (SoG) basin of the Salish Sea (Fig. 1). Most of these juveniles migrate northwards through the SoG into the tidally mixed Discovery Islands (DI) and Johnstone Strait (JS) before reaching the productive Queen Charlotte Strait (QCS) (McKinnell et al., 2014). Thomson et al. (2012) hypothesized that poor foraging conditions for juvenile salmon in the SoG and QCS before entering the Gulf of Alaska were responsible for high mortality and low adult returns for that year class. In the Trophic Gauntlet Hypothesis (TGH), McKinnell et al. (2014) went further to propose that these anomalous conditions in QCS / SoG were amplified by consistently poor foraging conditions in JS, where strong tidal mixing causes light limited primary production and low zooplankton biomass. Subsequent research has demonstrated that tidally mixed waters in JS are indeed characterized by poor foraging conditions, with a prevalence of small zooplankton (< 2 mm; Mahara et al. (2021)), corresponding with low foraging success for migrating juvenile sockeye salmon (James et al., 2020). Low insulin-like growth factor 1 (IGF-1) levels in JS compared to higher levels in the northern Strait of Georgia (NSoG), QCS and Queen Charlotte Sound supports an impact of these foraging conditions on juvenile salmon growth (Ferriss et al., 2014; Journey et al., 2018).

**Figure 1.**
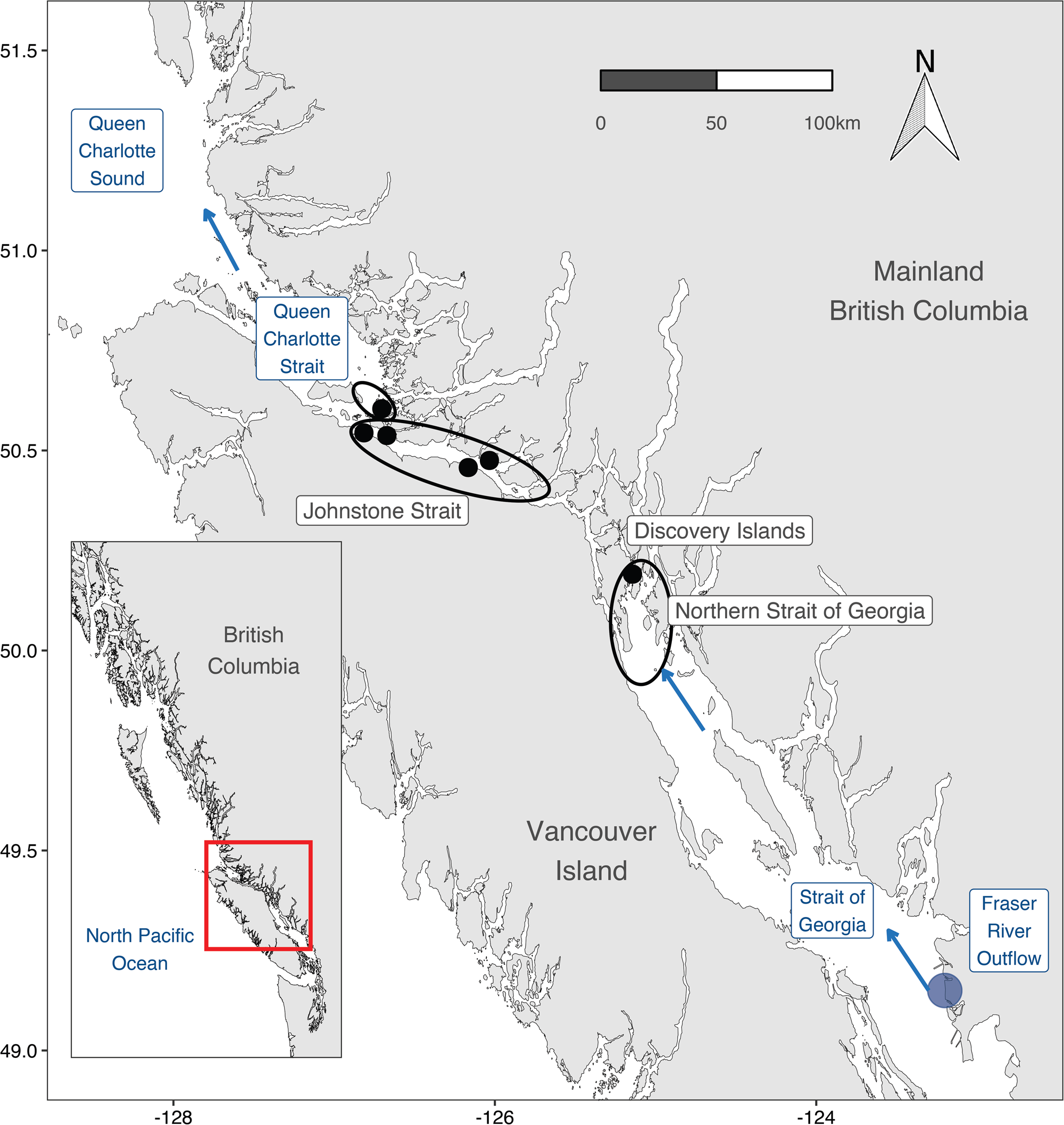
Map of sampling locations (black points) for juvenile salmon. Encircled points indicate the sampling regions, blue point indicates Fraser River outflow, and blue arrows indicate the primary Fraser sockeye salmon migration route. Inset map shows the location of the study area on the British Columbia coast. Figure created with R package ‘PBSmapping’ (Schnute et al., 2021).

Relating growth proxies to environmental conditions needs to consider the integration time of the proxy used. IGF-1 is a biochemical proxy of growth that has been shown to integrate feeding conditions experienced over 4-14 days (Beckman et al., 2004), relatively long when assessing the response of fast migrating organisms over short distances (Duguid et al., 2018; Gabillard et al., 2006; Pierce et al., 2005). The need for precise handling when sampling in the field, and the difficulty in sampling from individuals smaller than 120 mm also make the IGF-1 method difficult to routinely apply in juvenile salmon research (Duguid et al., 2018; Ferriss et al., 2014). Another growth proxy commonly used in fish ecology and fishery research is RNA/DNA ratio (Clemmesen and Doan, 1996). The underlying principle of using the ratio of RNA to DNA is that the concentration of DNA per cell is constant, whereas the RNA concentration per cell varies with its anabolic activity. The RNA/DNA ratio integrates feeding history over a period of 1-5 days (Buckley et al., 1999; Clemmesen and Doan, 1996; Wright and Martin, 1985), which makes it useful for assessing fish response to rapid spatial changes in coastal migration habitat. RNA/DNA can be effectively sampled irrespective of individual size and can be sampled from individuals frozen in the field without any degradation.

Another variable that needs to be considered when assessing fish growth response is foraging conditions. Prey quantity is often considered one of the main drivers of growth, survival, and abundance of marine organisms (Bilton and Robins, 2011), however, prey quality also plays a critical role (Litz et al., 2017). Fatty acids (FA), and essential fatty acids (EFA) in particular, are useful indicators of both food quality and the nutritional condition of a fish (Tocher, 2003). EFAs are not synthesised *de novo* by vertebrates in sufficient quantities to meet their physiological demands (Xu et al., 2018). Rather, EFAs are primarily synthesised by phytoplankton and subsequently reach consumers through trophic transfer (Dalsgaard et al., 2003). Marine carnivorous fish, such as sockeye salmon, have only a limited ability to convert EFAs into important long-chained PUFAs such as docosahexaenoic acid (DHA; 22:6n-3), eicosapentaenoic acid (EPA; 20:5n-3), and therefore must be obtained from their diets (NRC, 2011). Studies have shown that DHA and EPA concentrations measured in fish muscle tissue reflect the FA composition of their diets (Jin et al., 2017; Xu et al., 2020). Total amounts of DHA and EPA are known to affect the development, growth, and survival of juvenile marine fish (Mourente et al., 1991) and are of major importance during periods of rapid growth (Jin et al., 2017) (Table 1). Low levels of DHA and EPA can lead to reduced growth, maldevelopment, reduced reproduction, and ultimately decreased survival (Copeman and Parrish, 2002) (Table 1). Stored FAs, especially DHA and EPA, are also important when fish experience times of food limitation and starvation, for energy mobilization, hyperphagia, and for compensatory growth.

**Table 1.** Summary table of fatty acid groups, names and function.

| Fatty acid group | Fatty acid name | Fatty Acid Abbreviation | Fatty acid function | Reference |
| --- | --- | --- | --- | --- |
| **Highly unsaturated fatty acids** | docosahexaenoic acid;<br>eicosapentaenoic acid;<br>arachidonic acid | DHA; C22:6n-3<br>EPA; C20:5n-3<br>ARA; C20:4n-6 | Cell membranes, pigmentation, eye development, brain development, gene expression, neural cell development, Prostaglandin production | Parrish (2009) |
| **Saturated Fatty Acids (SFA)** |  |  | SFAs can be mobilized for ATP production | Wen et al. (2006) |
| **Monounsaturated Fatty Acids (MUFA)** |  |  | MUFAs can be mobilized for ATP production | Wen et al. (2006) |
| **Polyunsaturated Fatty Acids (PUFA)** |  |  | PUFAs can be mobilized for ATP production<br>Growth, survival, metamorphosis, pigmentation | Wen et al. (2006)<br>Copeman et al. (2002) |
|  |  | C16:0 (SFA),<br>C16:1 (MUFA),<br>C18:1n-9 *cis* (MUFA),<br>C18:2n-6 *cis* (PUFA) | Biomarker for starvation in fish | Torstensen et al. (2000) |

Experimental studies have shown that food quality changes, especially micronutrient limitation, and associated fatty acid changes lead to reduced RNA/DNA as lipid and protein metabolism is compromised when supply is limited (Boersma et al., 2008). The fatty acid composition integrates the feeding history over a period of approximately one week in juvenile Chinook salmon (Garzke et al. in rev), and using RNA/DNA and fatty acids in combination therefore provides a useful approach to identify changes in nutritional and physiological condition in organisms (Clemmesen, 1994; John et al., 2001; Malzahn et al., 2007; Paulsen et al., 2014; Xu et al., 2009)

In this study, we examined the response of juvenile sockeye salmon RNA/DNA, fatty acids and body mass index as they migrated through complex oceanographic habitats in coastal British Columbia, Canada. Specifically, our study area spanned the trophic gauntlet region (McKinnell et al., 2014) allowing assessment of the fish nutritional condition when migrating through JS, a region previously demonstrated to provide food poor conditions.

### Materials and Methods

#### Ethics Statement

This study did not involve endangered or protected species and was carried out under the guidelines of the Canadian Council on Animal Care. Juvenile salmon were collected under DFO license numbers ‘XR 42 2015’ and ‘XR 92 2016’ with approval from UBC’s Animal Care Committee (Protocol A19-0025). The study was carried out in compliance with the ARRIVE guidelines 2.0 (www.arriveguidelines.org).

#### Study location

Juvenile sockeye salmon were sampled at 6 locations between the NSoG and QCS (Fig. 1) in 2015 (6 May - 7 July) and 2016 (14 May - 6 July), in three regions along their outmigration route (Table S1). From south to north: the Discovery Islands (DI), JS, and southern QCS (Fig. 1). Based on oceanographic sampling (Dosser et al., 2021), the DI have water properties of the northern SoG and JS has water properties of QCS. The JS sites were all located within JS proper, where water depth was > 300 m. The QCS site was located in water <150 m deep to the north of the JS sill. Assuming RNA/DNA turnover rates of 1-5 days, the RNA/DNA ratios of fish collected in the DI were expected to predominantly reflect their migration experience through the northern SoG. Given this and the NSoG water properties, we henceforth refer to the DI as NSoG. Based on RNA/DNA replacement rates, JS fish would reflect conditions in the DIJS, and QCS fish would reflect the transition from JS to QCS. According to tagging data, the travel rates of juvenile sockeye between these regions are < 2 days for NSoG to DI, ca. 7 days from DI to JS, and ca. 4 days from JS to QCS (Furey et al., 2015; Johnston, 2020; Rechisky et al., 2018; Welch et al., 2011).

#### Juvenile Salmon Sampling

Fish were collected from a 6 m motorized vessel at distances of 5 - 60 m from shore, using a purse seine designed for manual deployment and retrieval (bunt: 27 × 9 m with 13 mm mesh; tow 46 × 9 m with 76 mm mesh) (Hunt et al., 2018). Captured fish were initially held alongside the vessel in a submerged portion of the seine’s bunt, allowing fish to swim without contacting the net and minimizing stress. Individual fish were transferred from the net by capturing them using a seawater-filled plastic jug with the end cut off. The fish were euthanized with MS-222 solution (250 mg L^-1^) in a 532 mL plastic bag with a unique identifier and frozen using liquid nitrogen vapour in a dry-shipper (MVE Doble-47). Individuals were stored at -80°C until further dissection in the lab. A total of 1,233 juvenile sockeye were collected across all stations and years (weight: min. 2.8 g – 32.8 g (mean: 12.66 g, median: 11.5 g); length: min. 63 mm – 145 mm (mean: 106.43 mm, median: 106 mm)), and a maximum of 20 individuals at each seine event. Genetic stock identification showed that 98% of the 2015 and 100% of 2016 were Fraser River salmon (Table S4).

#### Laboratory analysis

Five sockeye salmon were randomly selected from the total caught and were retained from each purse seine for a given site and day for further analysis of RNA/DNA and fatty acids (n=187). Before dissection fish were measured and weighed (mean weight = 9.07 g; range 1.27 – 38.85 g). White muscle tissue was collected from close to the dorsal fin for RNA/DNA analysis, and 400-600 mg of muscle tissue from close proximity to the dorsal fin for fatty acid analysis. Tissue samples were then returned to a -80 °C freezer until further analysis. In total, 187 juvenile salmon were processed across all regions and both years for fatty acids composition and RNA/DNA ratio (2015: n=87: NSoG (n=16), JS (n=34), QCS (n=37); 2016: n=100: NSoG (n=22), JS (n=37), QCS (n=41)).

#### RNA/DNA analysis

RNA/DNA was measured in crude muscle tissue homogenates using the non-specific, nucleic acid intercalating fluorescence dye RiboGreen (Gorokhova and Kyle, 2002). Nucleic acid quantification from white-muscle tissues followed a modified protocol by Caldarone et al. (2006) and Clemmesen et al. (1993). For RNA and DNA extraction, 1% N-lauroylsarcosine (VWR, Cat. No. VWRV0719-500G) was added to the samples, followed by homogenization with glass beads (VWR Cat. No. 12621-154), shaken for 30 min in a Retsch cell mill. Samples were centrifuged for 10 min at 14,000 x *g* and the supernatant diluted to match the RiboGreen saturation range (1:20, 1:30, 1:50, 1:100 and 1:200). Triplicate aliquots from the diluted sample were added to a black, flat-bottom 96-well microtiter plate. Standard solutions of rRNA (*Eschericia coli* 16S and 23S rRNA, Quant-iT RiboGreen RNA assay Kit Thermo Fisher Scientific Cat. No. R11490) and DNA (calf thymus, VWR, Supp. No. MB-102-01100) were diluted to final concentrations between 0 µg/mL and 3 µg/mL. 1x RiboGreen was added to each well and incubated for 5 min. Fluorescence was measured on a VarioSkan Flash Microplate Reader (ThermoFisher Scientific; excitation = 500 nm, emission= 525 nm), with the signal representing the fluorescence of RNA and DNA combined. Ribonuclease A (bovine pancreas, Alpha Aesar, Cat. No. CAAAJ62232-EX3) was added to each well and incubated at 37°C for 30 min. Fluorescence of the digested samples were measured to determine the DNA only. The RNA concentration was calculated by subtracting the DNA fluorescence and DNA+RNA fluorescence. The RNA/DNA was calculated for each well, and samples where the coefficient of variation (CV) for either RNA or DNA concentration in triplicate wells exceeded 15% were excluded from further analysis following.

#### Fatty acid analysis

Fatty acid analyses were performed following a slightly modified version of the protocol described in Puttick et al. (2009) using a one-step fatty acid methyl ester (FAME) method. Wet weights of all muscle samples were measured, freeze-dried, and dry weight measured for calculations of moisture content. FAMEs were obtained by reaction in a solution of 2.0 mL of 3 N HCl in CH_3_OH (Sigma-Aldrich cat. #90964-500ML). Nonadecanoic acid (C19:0) was used as an internal standard (Abdulkadir and Tsuchiya, 2008). FAMEs were analysed with a gas chromatograph (Scion 436-GC, Scion Instruments Canada, Edmonton, Alberta, Canada) using a flame ionization detector (FID). FAMEs were separated on a 50 m column (Agilent J&W CP-Sil 88 for FAME, Santa Clara, California, USA) using hydrogen as the carrier gas. Peaks were identified against an external standard (GLC 455 and GLC 37 Nu-chek Prep, Inc., Elysian, Minnesota, USA. For the purpose of this analysis the following fatty acid grouping were used: total fatty acids (TFA), saturated fatty acids (SFA), monounsaturated fatty acids (MUFA), polyunsaturated fatty acids (PUFA), EPA and DHA.

#### Oceanographic sampling

CTD profiles were collected every one to two weeks to characterise the regional oceanographic conditions. CTD stations were in close proximity (0-2 km) to the salmon sampling stations (Fig. S1). An RBR maestro or a SeaBird 19plus V2 CTD was used (Halverson et al., 2017). For the purpose of this study, we present temperature (°C) and salinity data integrated over the upper 10 m of the water column, where salmon were captured.

#### Zooplankton sampling

Zooplankton were collected approximately every fortnight in the NSoG, and every 4-7 days in JS and QCS (Fig. 1). A Bongo net (250 µm mesh size, mouth diameter 0.5 m) was deployed vertically from near-bottom or a maximum of 300 m depth to the surface. Filtered water volume during each tow was measured using a General Oceanics mechanical flowmeter. Collected zooplankton were preserved in 5% buffered formaldehyde-seawater solution. Zooplankton were identified to the lowest taxonomic level possible, counted, and taxon specific biomass was estimated by multiplying the mean dry weight (DW) of each species and stage by their abundance (Mahara et al., 2019). Zooplankton abundance and biomass data were integrated over the water column and presented as ind. m^-2^ and mg dry weight (DW) m^-2^. Since the zooplankton samples were integrated over 0-300 m depth, they overestimate the available zooplankton biomass and abundance for juvenile salmon, given that they likely forage in the top 30 m. Therefore, for the purpose of this study we only included zooplankton species that had been demonstrated to be consumed by the juvenile sockeye salmon in 2015 and 2016 (see species list Table S2, based on James (2019)).

#### Statistical analyses

Juvenile sockeye salmon length-weight residuals were calculated for all collected fish that were used for RNA/DNA and fatty acid analyses from both 2015 and 2016 (n=187 out of n=1,233) based on ln(weight) to ln(fork length) residuals from ordinary least squares linear regression (Reist, 2011). Positive length-weight residuals indicate that the salmon was heavier than expected for their fork length. We used generalized linear mixed models (lme4 (version 1.1-27.1) and lmerTest (version 3.1-3) in R) to examine the main and interactive effects of year and region (both categorical fixed factors on fish condition metrics (RNA/DNA, TFA, SFA, MUFA, PUFA, EPA+DHA, and length-weight residuals), further seine id was fit as a random effect on the intercept. Prior to this analysis, normal distribution was tested and if the data were not normally distributed the probability distribution was set for log normal (temperature and zooplankton biomass, and RNA/DNA). Differences were deemed significant where p < 0.05.

## Results

### Juvenile salmon condition

Fish length-weight residuals differed significantly between years (p < 0.05, Table 2) but not between JS and NSoG and JS and QCS (JS-NSoG: p=0.283; JS-QCS: p=0.569, Table 2). Overall, juvenile salmon were significantly heavier than expected based on their length in 2015, whereas in 2016 they were lighter than expected (Fig. 2A).

**Table 2.**
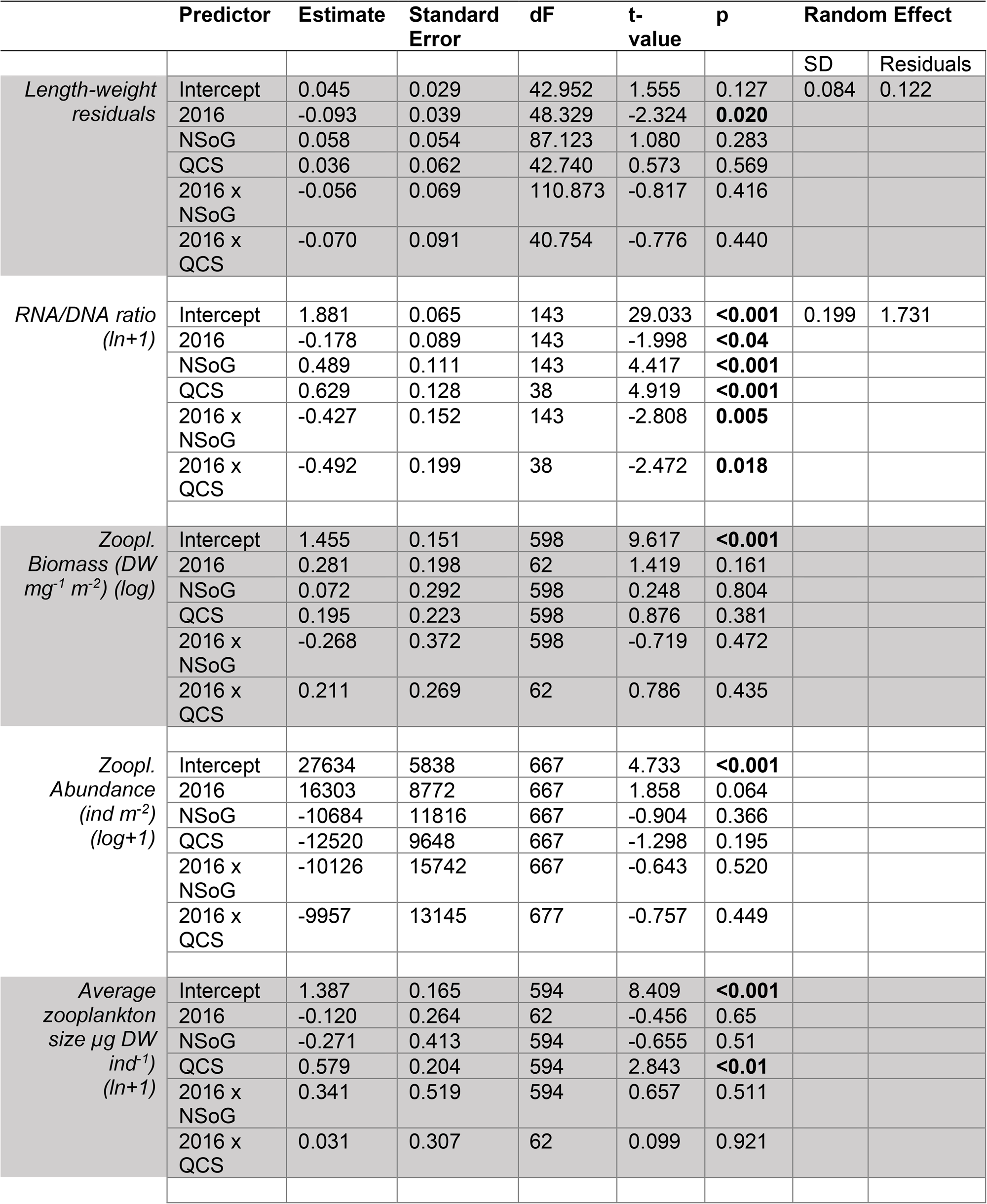
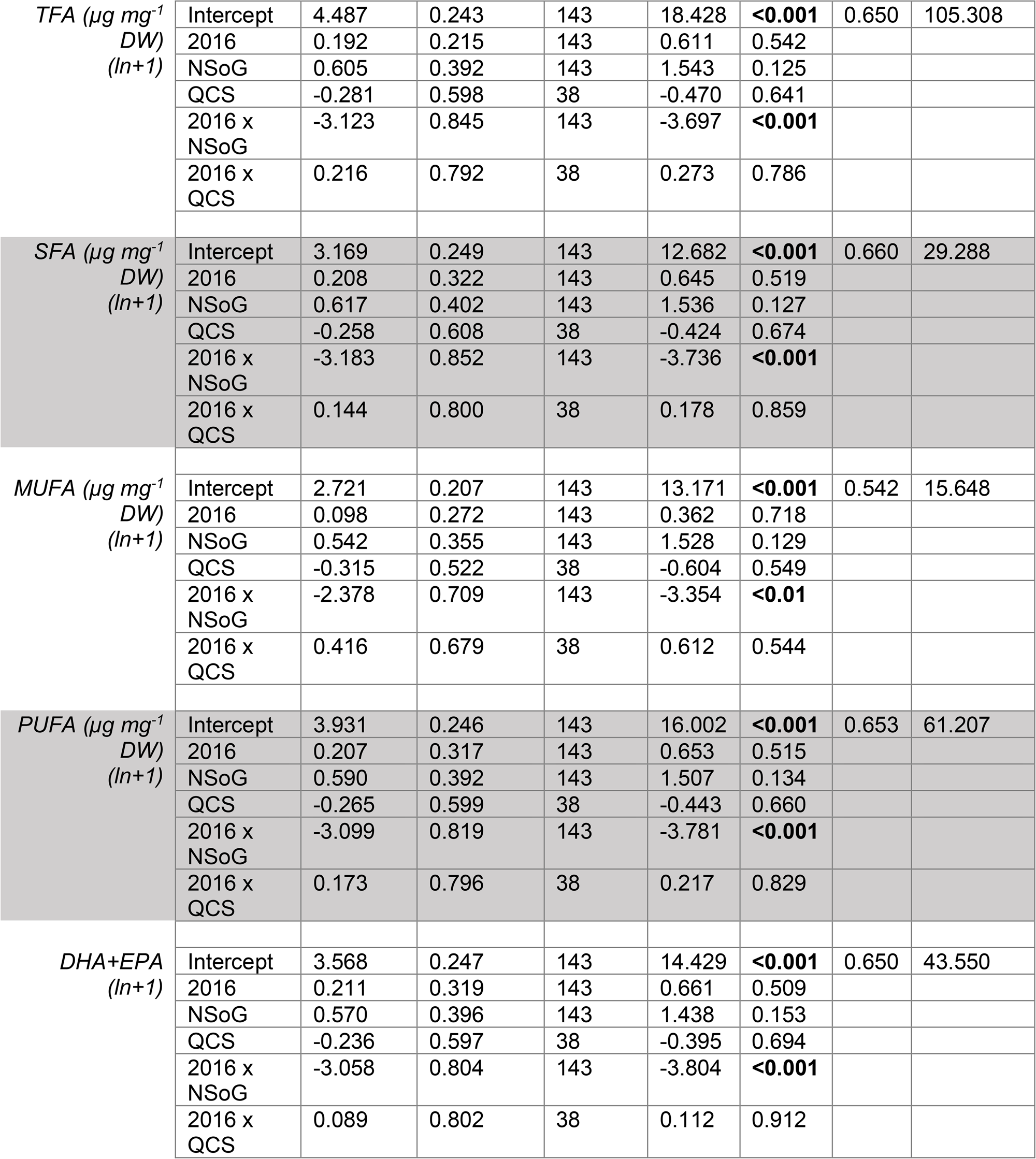
Results of generalized linear mixed models explaining the effects of region, year, and the interaction (x) of region and year. Values in bold are significant at p < 0.05.

**Figure 2.**
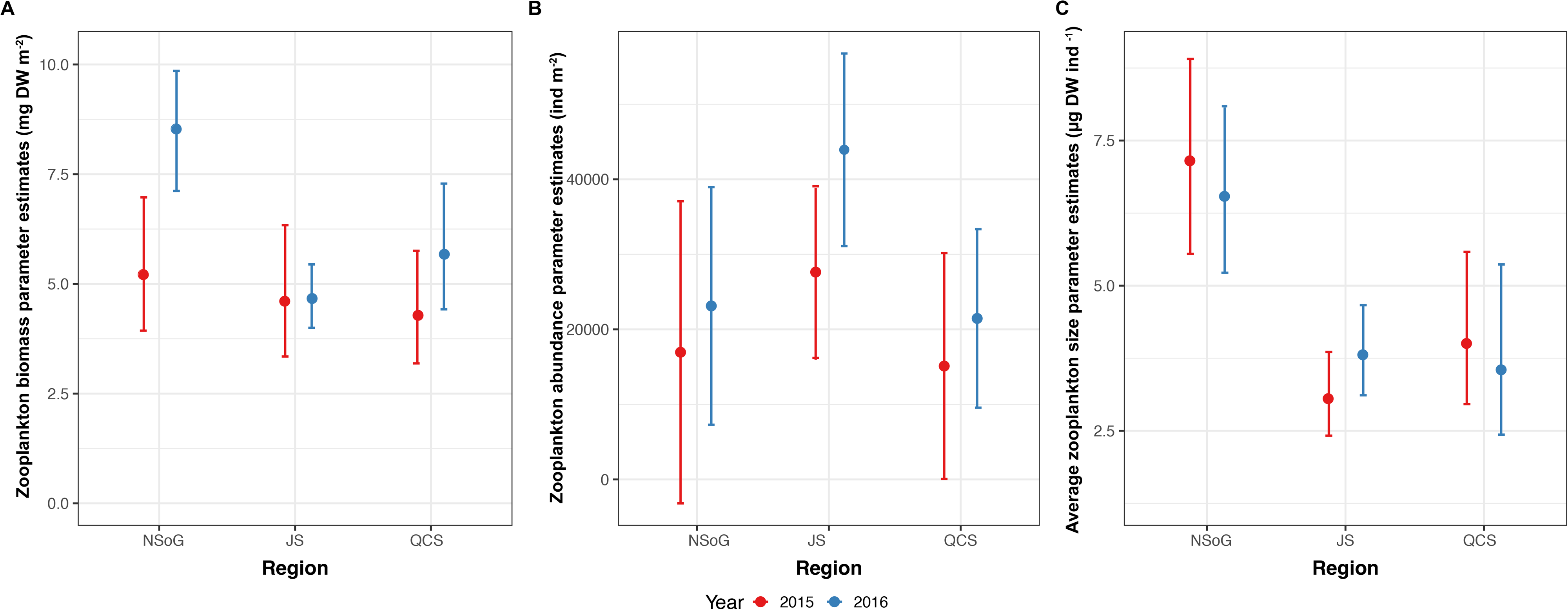
Annual and regional model parameter estimates of A) length-weight residuals of fish sampled for RNA/DNA (black; n=162), and B) RNA/DNA ratio. Error bars denote ± 95% Confidence Intervals. NSoG = northern Strait of Georgia; JS = Johnstone Strait; QCS = Queen Charlotte Strait; 2015 is indicated by red and 2016 by blue.

Juvenile salmon RNA/DNA differed significantly between years, regions, and due to the interaction of year and region (Table 2). RND/DNA ratios were significantly higher in 2015 than 2016, and independent of year, NSoG and QCS juvenile salmon had significantly higher RNA/DNA ratios than fish from JS (Table 2, Fig. 2B). In 2015, both NSoG and QCS juvenile salmon had significantly higher RNA/DNA ratios than 2016 (Table 2, Fig. 2B). In 2015, juvenile sockeye salmon entered the NSoG with high RNA/DNA ratios, which significantly decreased in JS, and then increased again in QCS (Fig. 2B). In 2016, RNA/DNA ratios were low across all stations, as low or lower than values in JS in 2015 (Fig. 2B). There was no significant correlation between fish RNA/DNA ratio and fish size (R = 0.07, p = 0.34, df = 163, Table S3) or between RNA/DNA and sea surface temperature (R = -0.11, p = 0.12, df = 185).

### Zooplankton

There was no significant difference between years or regions for either zooplankton biomass or zooplankton abundance (Table 2, Fig. 3). Average individual zooplankton size (Fig. 3C), calculated as total prey species biomass divided by total prey species abundance, was significantly higher in QCS (p < 0.01), with the smallest average zooplankton size in NSoG and JS (2.52µg DW ind^-1^ ± 6.10 SD) and largest size in QCS (5.74 µg DW ind^-1^ ± 11.13 SD). There was no difference in average zooplankton size between NSoG and JS (p = 0.51).

**Figure 3.**
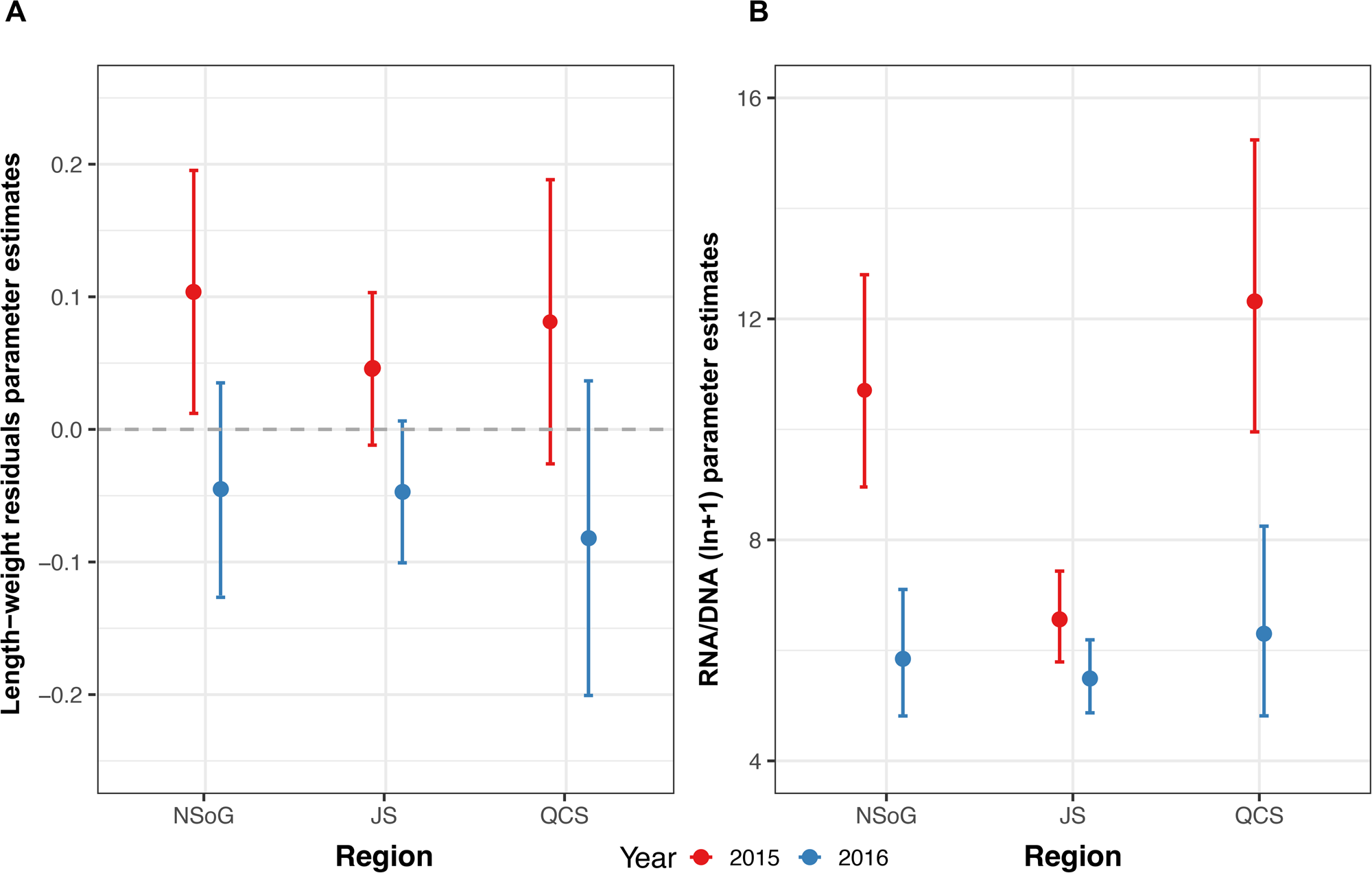
Annual and regional model parameter estimates of A) biomass, B) abundance of zooplankton > 1 mm in length, and C) average zooplankton size (µg DW ind^-1^). Error bars denote ± 95% Confidence Intervals. NSoG = northern Strait of Georgia; JS = Johnstone Strait; QCS = Queen Charlotte Strait; 2015 is indicated by red and 2016 by blue.

### Fatty acid profiles of juvenile salmon

Juvenile salmon TFA, SFA, MUFA, PUFA and DHA+EPA had similar regional and annual trends (Fig. 4A-E). There was a significant interaction between region and year for all fatty acid metrics (Table 2). Fatty acid concentrations were significantly higher in the NSoG in 2015 compared to 2016 (Fig. 4A-E). In 2015, juvenile salmon fatty acid levels were highest in the NSoG, decreased markedly in JS, and then slightly increased in QCS (Fig. 4A-E). In 2016, FA levels increased slightly from the NSoG to JS, and increased further in QCS to reach levels similar to those in 2015 (Fig. 4 A-E).

**Figure 4.**
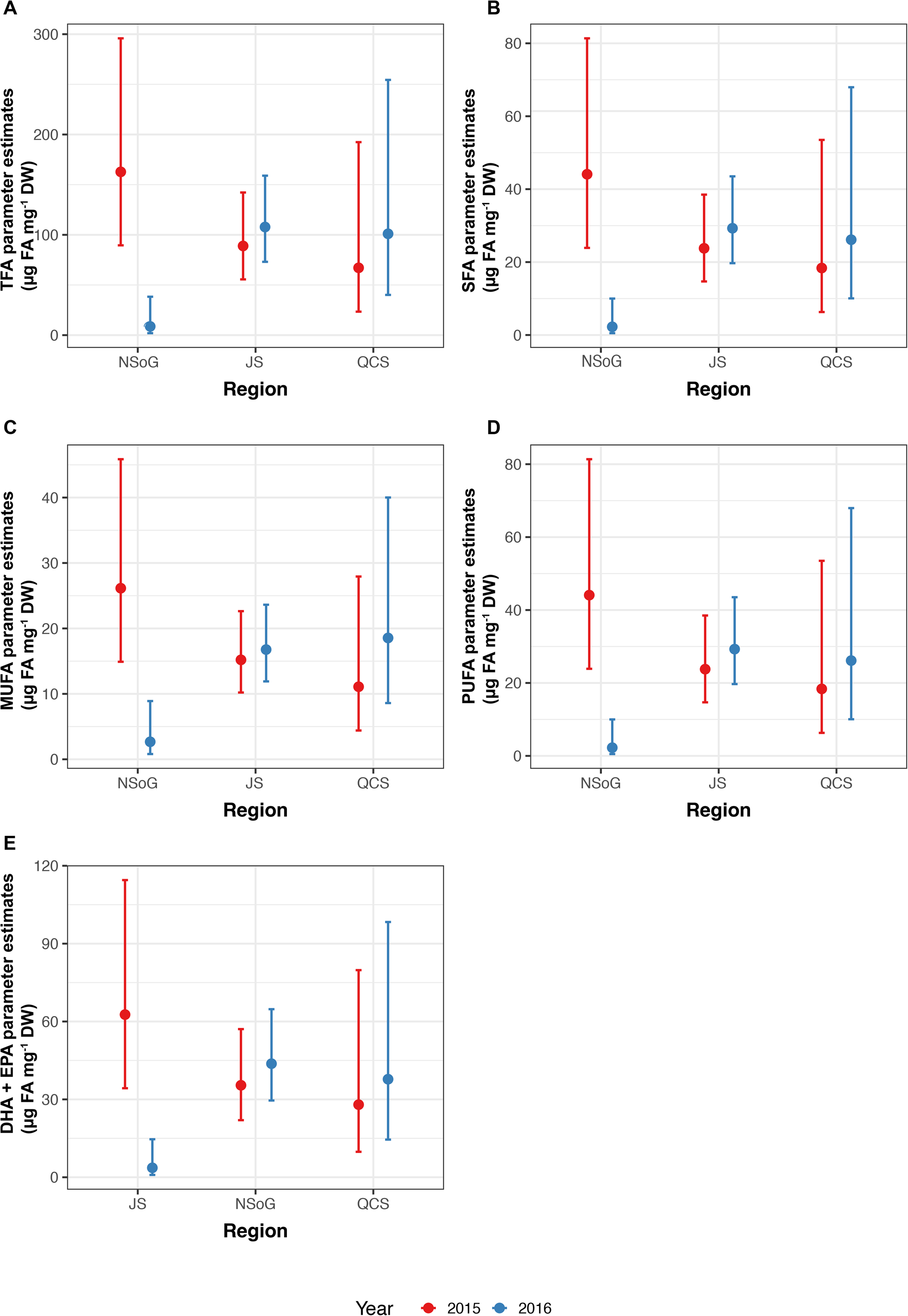
Annual and regional model parameter estimates of fatty acid concentrations (µg FA per mg dry weight of white muscle tissue) A) total fatty acids (TFA), B) saturated fatty acids (SFA), C) monounsaturated fatty acids (MUFA), D) polyunsaturated fatty acids (PUFA), and E) DHA + EPA (docosahexaenoic acid + eicosapentaenoic acid). Error bars denote ± 95% Confidence Intervals. NSoG = northern Strait of Georgia; JS = Johnstone Strait; QCS = Queen Charlotte Strait; 2015 is indicated by red and 2016 by blue.

## Discussion

Juvenile salmon encounter diverse ocean conditions during their early marine life. In this study, we compared the body condition of juvenile salmon (RNA/DNA) to their nutritional condition (fatty acid profiles) and prey availability (zooplankton biomass) across an ∼ 120 km section of their migration route in coastal British Columbia, in 2015 and 2016. Both body and nutritional condition of juvenile sockeye salmon showed short term changes that differed between years but were not related to zooplankton biomass, abundance, average size, or species composition. Our RNA/DNA data showed that the recovery of juvenile salmon in Queen Charlotte Strait (QCS) after a period of starvation in Johnstone Strait (JS) might depend on their condition prior to entering JS. Furthermore, our study provides evidence that the Trophic Gauntlet Hypothesis should be extended to include food quality (e.g., fatty acid content), not just food quantity. Below we discuss the role of fish condition in areas with coastal oceanographic variation, and the cumulative effect of fish encountering a mosaic of habitats during migration.

As outlined in the introduction, the majority of outmigrating juvenile Fraser River sockeye salmon travel northwards through the Strait of Georgia (SoG) before entering a section of tidally mixed channels in the Discovery Islands (DI) and JS (Furey et al., 2015). In the SoG, they typically encounter surface waters that are relatively warm, fresh, and stratified, correlated with high phytoplankton biomass, high zooplankton biomass and high mean zooplankton size (James et al., 2020; Mahara et al., 2021). Ocean conditions become cooler and saltier as the fish move north into the DI, JS, and QCS. This change in ocean conditions is driven by the deep tidal mixing that brings cold and salty deep water to the surface (McKinnell et al., 2014). Phytoplankton biomass is low through the tidally mixed northern DI and JS, leading to both low zooplankton biomass and smaller zooplankton size. This pattern is, however, reversed in QCS.

In the present study, juvenile salmon sampled in 2015 had RNA/DNA ratios which supported the Trophic Gauntlet Hypothesis (McKinnell et al., 2014), with high fish condition in the NSoG and QCS, and low condition in JS. The link between RNA/DNA and feeding success is supported by juvenile sockeye salmon stomach fullness analysis across the study area in 2015, which found high stomach fullness in the NSoG (Gut Fullness Index (GFI) = 1-2 %), low fullness in DI (GFI = 0.2 %), low fullness in JS (GFI = 0.48 %), and full stomachs in QCS (GFI = 3.5%) (James et al., 2020). Poor foraging conditions for juvenile sockeye salmon, with GFI < 1, were also observed in 2016 (James, 2019). Length-weight residuals showed a weaker response over the migration route, likely reflecting their longer integration times and highlighting the importance of more sensitive growth proxies, with faster turnover, to understand the response of fish to small scale regional changes in environmental conditions. Notably, RNA/DNA ratios and length-weight residuals showed a significantly different pattern in 2016. RNA/DNA ratios and length-weight residuals were significantly lower in NSoG juvenile salmon compared to 2015, indicating that feeding conditions in the NSoG were poorer in 2016 than 2015. In 2016, RNA/DNA ratios remained low during the migration through JS and increased only slightly in QCS. The lower condition in 2016 occurred despite similar zooplankton biomass, abundance, and average size to 2015, indicating that another factor was contributing to juvenile salmon nutritional and body condition.

There was no correlation between RNA/DNA and SST, indicating that temperature did not lead to the interannual difference in body condition. However, there were significant differences in juvenile sockeye salmon fatty acid profiles both between regions and years. Stored lipids and fatty acids in fish tissues are important to mobilize energy from muscle, liver, and viscera by FA β-oxidation processes during periods of starvation (Halver and Hardy, 1989). During starvation, saturated FA (SFA) are the first FAs to be mobilized for ATP production, followed by monounsaturated FA (MUFA) and then polyunsaturated FA (PUFA) (Wen et al., 2006). In 2015, we detected a decrease of TFA, SFA, MUFA, and PUFA along the migration corridor, with lowest concentrations in JS, indicating that fish had oxidized FAs stored in white muscle tissue for ATP production. In 2016, juvenile salmon sampled in the NSoG had significantly lower concentrations of TFA, SFA, MUFA, and PUFA than 2015, indicating that the nutritional condition of salmon was already very low when they reached the NSoG, with less FAs stored in white muscle tissue as an energy reserve for starvation periods.

As FAs have a response time of a minimum of one week, the condition of juvenile salmon sampled in the NSoG in this study must have been in response to the preceding conditions in the SOG. Differences between years in body and nutritional condition may have resulted because either food quantity or food quality were low in the SoG in 2016, leading to depletion of FA reserves needed to meet the energy demands of migration. Zooplankton biomass in the study area was not significantly different between 2015 and 2016 (this study; Mahara et al. (2019)) and was significantly higher in the central SoG in 2016 (Perry et al., 2021). This suggests that lower food quality (e.g., DHA + EPA) in 2016 was the most likely driver of lower fish FA content and RNA/DNA ratios in that year. As outlined in the introduction, low DHA and EPA availability in prey have important implications for fish condition, including reduced growth rates, visual maldevelopment with negative effects on hunting efficiency, an increase in FAs that are considered as pro-inflammatory (Table 1), and reduced capacity for compensatory growth, i.e., accelerated growth that follows a period of limited growth once non-limiting conditions are encountered (Ballantyne et al., 2003; Bou et al., 2017).

Fish obtain DHA and EPA through their diet. In the case of zooplankton prey, variations in DHA and EPA can occur because of shifts in composition, with different species having different nutritional quality and fatty acid content (Hiltunen et al., 2021), or shifts in the nutrition of phytoplankton that the zooplankton feed on (Costalago et al., 2020; El-Sabaawi et al., 2009). (2009). There is substantial variability in FA content among zooplankton species in the Salish Sea (El-Sabaawi et al., 2009; Hiltunen et al., 2021). However, analysis of zooplankton communities has shown that species composition in the NSoG did not differ between 2015 and 2016 (Mahara et al., 2019), indicating that this was not an important factor in zooplankton prey quality. It should be noted that similarly, there was no interannual difference in zooplankton community composition in the DI, JS, or QCS either (Mahara et al., 2021). With regard to zooplankton’s phytoplankton prey, no major differences were observed in phytoplankton composition in the NSoG between 2015 and 2016 (Belluz et al., 2021). However, spring bloom timing differed substantially between years: in 2015 an unusually early spring bloom (February 24^th^) occurred, while in 2016 the bloom timing was typical (April 1^st^) with maximum biomass of 12.40 mg Chl *a* m^-3^ and 11.53 mg Chl *a* m^-3^, respectively (Belluz et al., 2021; Mahara et al., 2019). It is possible that the later phytoplankton bloom in 2016 delayed the development of the zooplankton community and onset of lipid accumulation in the later life stages (e.g., Mayzaud et al. (1999))

The better body and nutritional condition of juvenile salmon leaving the NSoG in 2015 appeared to effect recovery after leaving JS. In 2015, juvenile salmon RNA/DNA ratios in QCS showed compensatory growth, almost doubling after passing JS where RNA/DNA ratios were low and FA concentrations declined due to necessary energy mobilization. In contrast, in 2016 when juvenile salmon already had low condition and depleted energy reserves in the NSoG, no compensatory growth was observed in QCS. Studies with Atlantic salmon (*Salmo salar*) have shown that compensatory growth success depends on the quality of food available, particularly requiring high lipid content (Johansen et al., 2001). Turchini et al. (2007) proposed the term ‘lipo-compensatory growth’ after showing that Murray cod were able to compensate for starvation and refuel fatty acid storage when fed with high fatty acid diets. This supports the importance of high-quality food for enduring periods of poor foraging conditions.

Protracted periods of poor foraging may not only slow down recovery but may also have long-term impacts on the physiology of the fish (Nikki et al., 2004). Periods with very low food biomass longer than 3-4 weeks can decrease digestive enzyme activity, inhibiting the ability of fish to sufficiently digest food when available again, resulting in overall lower body size and higher mortality (Abolfathi et al., 2012). This is highly relevant to juvenile salmon migrating through the BC coastal ocean. The migration of juvenile sockeye salmon through the DI and JS takes approximately 14 days, while time spent in the SoG is estimated to be between 30 and 50 days (Preikshot et al., 2012). Thus, as suggested by McKinnell et al (2014), anomalously poor foraging conditions in the SoG and/or QCS, coupled with typically poor foraging conditions in the DI and JS, may have an additive effect that impacts early marine survival of juvenile salmon. Our study goes further, providing evidence that the Trophic Gauntlet Hypothesis should be extended to food quality as a critical factor in juvenile salmon growth and survival.

## Conclusions

This study demonstrated that both body and nutritional condition varied between regions and years across a dynamic stretch of their British Columbia migration route. Our data supported JS as a region of poor foraging success that significantly affected fish health. The impact of JS appeared to be modulated by the body condition with which the fish entered the region. In 2015, when juvenile sockeye salmon entered JS in good body and nutritional condition, fish were able to use stored resources to metabolize stored energy when traversing JS to QCS where high prey abundance enabled compensatory growth. In 2016, when juvenile sockeye salmon entered JS in poor body and nutritional condition, compensatory growth was not detected in QCS. We hypothesize that lower zooplankton prey quality was responsible for the poor nutritional condition of juvenile salmon in the NSoG in 2016. Further, we propose that the Trophic Gauntlet Hypothesis should be extended to include to the nutritional quality of prey. The recovery of juvenile salmon from regions with naturally poor foraging conditions may depend on the availability and quality of prey in adjacent areas and be negatively affected by extreme climate events and climate change that reduce the quantity and quality of prey in historically productive areas.

## Acknowledgements

This research was conducted as part of the Hakai Institute’s Juvenile Salmon Program, funded by the Tula Foundation. J Garzke was supported by the Tula-Mitacs Canada Grants IT09911 and IT13677. Juvenile salmon were collected under DFO license number ‘XR 42 2015’ and ‘XR 92 2016’ with approval from UBC’s Animal Care Committee (Protocol A19-0025). We thank the field crews of the Hakai Institute’s Quadra Island research station and Salmon Coast Field Station for sample collection. We are grateful to have worked on the traditional, and ancestral lands of Da’naxda’xw/Awaetlala, We Wai Kum, We Wai Kai, Tlowitsis, Homalco (Xwemalhkwu), Kwakwaka’wakw, <u>K’ómoks</u>, Coast Salish peoples, and Klahoose Nations.

## Author contributions

**JG** - Laboratory and Statistical analysis, Visualization, Writing and Editing; **BPVH** – Project Conceptualization, Methodology, Writing and Editing; **SG** – Project Conceptualization, Methodology, Field program, Investigation, Editing; **IF, BTJ** - Project Conceptualization, Methodology, Field program, Investigation, Editing **MK** - Project Conceptualization, Methodology, Field program, Investigation, Editing; **NM** - Field program, Zooplankton Data, Editing; **EAP** - Project Conceptualization, Methodology, Editing; **LR** - Project Conceptualization, Methodology, Field program, Investigation, Editing.

## Additional Information

**Supplementary information** accompanies this paper

## Competing financial interest

The authors declare no competing financial interest

## Notes

### Competing Interest Statement

The authors have declared no competing interest.

### Summary of Updates

The statistical analyses has changed and accordingly the results section, discussion, and figures.

