## Supplementary material for "Dynamic coastal pelagic habitat drives rapid changes in growth and condition of juvenile sockeye salmon (*Oncorhynchus nerka*) during early marine migration": Suppl.

**Supplementary Materials**

**Table S1.** Sample distribution over the two field campaigns in 2015 and 2016

| | **Sampling site ID** | | | | | | |
|---|---|---|---|---|---|---|---|
| **Sampling date** | **NSoG D07** | **JS J03** | **JS J06** | **JS J07** | **JS J09** | **QCS J02** | **Total** |
| 2015-05-06 | 1 | 0 | 0 | 0 | 0 | 0 | 1 |
| 2015-05-16 | 3 | 0 | 0 | 0 | 0 | 0 | 3 |
| 2015-05-24 | 0 | 0 | 5 | 0 | 0 | 5 | 10 |
| 2015-05-26 | 2 | 5 | 0 | 0 | 4 | 0 | 11 |
| 2015-05-31 | 0 | 5 | 0 | 0 | 5 | 0 | 10 |
| 2015-06-01 | 0 | 0 | 0 | 0 | 0 | 2 | 2 |
| 2015-06-06 | 0 | 0 | 0 | 3 | 0 | 0 | 3 |
| 2015-06-07 | 5 | 0 | 0 | 0 | 3 | 0 | 8 |
| 2015-06-08 | 0 | 4 | 0 | 0 | 0 | 0 | 4 |
| 2015-06-09 | 0 | 0 | 0 | 0 | 0 | 4 | 4 |
| 2015-06-13 | 5 | 0 | 0 | 0 | 0 | 0 | 5 |
| 2015-06-14 | 0 | 0 | 0 | 3 | 0 | 0 | 3 |
| 2015-06-15 | 0 | 4 | 0 | 0 | 0 | 0 | 4 |
| 2015-06-16 | 0 | 0 | 0 | 0 | 6 | 3 | 9 |
| 2015-06-21 | 0 | 0 | 4 | 0 | 0 | 0 | 4 |
| 2015-06-23 | 0 | 1 | 0 | 0 | 5 | 0 | 6 |
| 2016-05-14 | 4 | 0 | 0 | 0 | 0 | 0 | 4 |
| 2016-05-21 | 0 | 0 | 0 | 3 | 0 | 0 | 3 |
| 2016-05-22 | 0 | 0 | 0 | 0 | 6 | 5 | 11 |
| 2016-05-26 | 3 | 0 | 0 | 0 | 0 | 0 | 3 |
| 2016-05-27 | 0 | 0 | 0 | 0 | 5 | 0 | 5 |
| 2016-05-28 | 0 | 0 | 0 | 0 | 0 | 5 | 5 |
| 2016-06-01 | 0 | 0 | 0 | 0 | 0 | 4 | 4 |
| 2016-06-02 | 0 | 0 | 0 | 0 | 3 | 0 | 3 |
| 2016-06-03 | 5 | 0 | 0 | 3 | 0 | 0 | 8 |
| 2016-06-04 | 0 | 5 | 5 | 0 | 0 | 0 | 10 |
| 2016-06-09 | 5 | 0 | 0 | 0 | 4 | 0 | 9 |
| 2016-06-10 | 0 | 5 | 0 | 0 | 0 | 0 | 5 |
| 2016-06-11 | 0 | 0 | 0 | 2 | 0 | 0 | 2 |
| 2016-06-16 | 5 | 0 | 0 | 0 | 0 | 0 | 5 |
| 2016-06-18 | 0 | 0 | 0 | 0 | 5 | 0 | 5 |
| 2016-06-19 | 0 | 0 | 5 | 0 | 0 | 0 | 5 |
| 2016-06-23 | 0 | 5 | 0 | 0 | 0 | 0 | 5 |
| 2016-06-25 | 0 | 0 | 4 | 0 | 0 | 0 | 4 |
| 2016-07-04 | 0 | 0 | 0 | 0 | 4 | 0 | 4 |
| **Total** | 38 | 34 | 23 | 14 | 50 | 28 | 187 |

**Table S3.** Mean fork lengths, standard deviations (SD), and 95% confidence intervals (CI) of sampled juvenile salmon during the field campaigns in 2015 and 2016 in the northern Strait of Georgia (NSoG), Johnstone Strait (JS), and Queen Charlotte Strait (QCS).

| **Year** | **Region** | **N** | **Mean fork length (mm)** | **SD** | **CI** |
|---|---|---|---|---|---|
| 2015 | QCS | 53 | 112.15 | 12.04 | 3.31 |
| 2015 | JS | 358 | 110.80 | 10.34 | 1.07 |
| 2015 | NSoG | 186 | 108.97 | 13.48 | 1.95 |
| 2016 | QCS | 69 | 102.72 | 8.45 | 2.02 |
| 2016 | JS | 429 | 104.51 | 12.59 | 1.19 |
| 2016 | NSoG | 138 | 97.26 | 10.24 | 1.72 |

**Table S4.** Genetic Stock IDs, following Beacham et al (2005), of all sampled juvenile fish (n=187). Northern Strait of Georgia (NSoG), Johnstone Strait (JS), and Queen Charlotte Strait (QCS). The percentage gives the highest probability of all available and tested genetic stock information.

| Sampling site | Lake | Region | Designatable Unit (DU) | 2015 | 2016 |
|---|---|---|---|---|---|
| **NSoG = D09** | Birkenhead Lake | Summer; Fraser River | DU12 | 0.95% | 0.00% |
| | Blackwater River | Early Summer; Fraser River | DU20 | 0.47% | 0.00% |
| | Chilko Lake | Summer; Fraser River | DU3 | 3.32% | 2.84% |
| | Chilko Lake -North | Summer; Fraser River | DU3 | 0.00% | 0.95% |
| | Gates Creek | Early Summer; Fraser River | DU3 | 0.47% | 0.00% |
| | Horsefly River | Summer; Fraser | DU1 | 0.00% | 0.95% |
| | Kynock | Early Stuart; Fraser River | DU16 | 0.47% | 0.00% |
| | Lake Adams | Late; Fraser River | DU20 | 2.37% | 2.84% |
| | Lake Shuswap | Late; Fraser River | DU18 | 0.00% | 3.32% |
| | Little Shuswap | Late; Fraser River | DU18 | 0.00% | 1.42% |
| | Middle Shuswap | Late; Fraser River | DU18 | 0.00% | 2.37% |
| | Mitchell River | Summer; Fraser River | DU18 | 1.42% | 0.00% |
| | Nahatlatch Lake | Early Summer; Fraser River | DU16 | 0.47% | 0.00% |
| | Pitt River | Early Summer; Fraser River | DU13 | 1.42% | 0.00% |
| | Quesnel Decept | Summer; Fraser River | DU15 | 0.47% | 0.00% |
| | Quesnel Horsefly | Summer; Fraser River | DU16 | 2.37% | 0.00% |
| | Quesnel Mitchell | Summer; Fraser River | DU16 | 2.37% | 0.00% |
| | Scotch Creek | Early Summer; Fraser River | DU16 | 0.47% | 0.47% |
| | Seymour River | Early Summer; Fraser River | DU18 | 0.47% | 0.00% |
| | Stellako River | Summer; Fraser River | DU19 | 1.42% | 0.00% |
| | Tachie River | Summer; Fraser River | DU7<br>DU21 | 0.47% | 0.00% |
| **QCS = J02** | Chilko Lake | Summer; Fraser River | DU3 | 2.84% | 0.00% |
| | Horsefly River | Summer; Fraser River | DU16 | 0.00% | 1.42% |
| | Lake Adams | Late; Fraser River | DU18 | 0.47% | 1.42% |
| | Lake Shuswap | Late; Fraser River | DU18 | 0.00% | 0.95% |
| | Mitchell River | Summer; Fraser River | DU16 | 0.95% | 0.00% |
| | Narrow Lake | Early Stuart; Fraser River | DU20 | 0.47% | 0.00% |
| | Pinchi Creek | Summer; Fraser River | DU21 | 0.95% | 0.00% |
| | Pitt River | Early Summer; Fraser River | DU15 | 0.47% | 0.00% |
| | Quesnel Horsefly | Summer; Fraser River | DU16 | 0.95% | 0.00% |
| | Quesnel Mitchell | Summer; Fraser River | DU16 | 0.47% | 0.00% |
| | Roaring Lake | Summer; Fraser River | DU16 | 0.47% | 0.00% |
| | Stellako River | Summer; Fraser River | DU7 | 0.00% | 0.47% |

| | | | | | |
|---|---|---|---|---|---|
| | Thompson North | Early Summer; Fraser River | DU11 | 0.95% | 0.00% |
| **JS = J03** | Birkenhead Lake | Summer; Fraser River | DU12 | 0.47% | 0.00% |
| | Chilko Lake | Summer; Fraser River | DU3 | 1.90% | 1.42% |
| | Chilko Lake South | Summer; Fraser River | DU3 | 0.47% | 0.00% |
| | Chilko Lake -North | Summer; Fraser River | DU3 | 0.00% | 0.47% |
| | Kynock | Early Stuart; Fraser River | DU20 | 0.00% | 0.47% |
| | Lake Adams | Late; Fraser River | DU18 | 0.47% | 0.47% |
| | Lake Shuswap | Late; Fraser River | DU18 | 0.00% | 1.90% |
| | Little Lake | Late; Fraser River | DU18 | 0.00% | 0.47% |
| | Middle Lake | Summer; Fraser River | DU18 | 0.47% | 0.00% |
| | Middle Shuswap | Late; Fraser River | DU18 | 0.00% | 0.47% |
| | Mitchell River | Summer; Fraser River | DU16 | 1.42% | 0.00% |
| | Pinchi Creek | Summer; Fraser River | DU21 | 0.47% | 0.00% |
| | Pitt River | Early Summer; Fraser River | DU15 | 0.95% | 0.00% |
| | Quesnel Horsefly | Summer; Fraser River | DU16 | 0.47% | 1.42% |
| | Upper Horsefly | Summer; Fraser River | DU16 | 0.47% | 0.00% |
| **JS = J06** | Chilko Lake | Summer; Fraser River | DU3 | 0.47% | 0.95% |
| | Chilko Lake south | Summer; Fraser River | DU3 | 0.47% | 0.00% |
| | Gates Creek | Early Summer; Fraser River | DU1 | 0.47% | 0.00% |
| | Horsefly River | Summer; Fraser River | DU16 | 0.00% | 1.90% |
| | Kuzkwa Creek | Summer; Fraser River | DU21 | 0.47% | 0.00% |
| | Lake Adams | Late; Fraser River | DU18 | 0.47% | 2.37% |
| | Lake Shuswap | Late; Fraser River | DU18 | 0.00% | 0.95% |
| | Mitchell River | Summer; Fraser River | DU16 | 0.47% | 0.00% |
| | Porter Creek | Early Stuart; Fraser River | DU20 | 0.95% | 0.00% |
| | Quesnel Horsefly | Summer; Fraser River | DU16 | 0.00% | 0.47% |
| | Seymour River | Early Summer; Fraser River | DU18 | 0.47% | 0.00% |
| **JS = J07** | Chilko Lake | Summer; Fraser River | D4 | 1.90% | 0.47% |
| | Lake Adams | Late; Fraser River | DU18 | 0.47% | 0.47% |
| | Pitt River | Early Summer; Fraser River | D15 | 0.47% | 0.00% |
| | Quesnel Horsefly | Summer; Fraser River | D16 | 0.00% | 0.95% |
| | Seymour River | Early Summer; Fraser River | DU18 | 0.00% | 0.95% |
| | Stellako River | Summer; Fraser River | DU7 | 0.00% | 0.95% |
| **JS = J09** | Birkenhead Lake | Summer; Fraser River | DU12 | 0.00% | 0.47% |
| | Chilko Lake | Summer; Fraser River | DU3 | 3.32% | 0.95% |
| | Chilko Lake south | Summer; Fraser River | DU3 | 1.90% | 0.00% |
| | Horsefly River | Summer; Fraser River | DU16 | 0.00% | 0.95% |
| | Lake Adams | Late; Fraser River | DU18 | 0.47% | 0.95% |
| | Lake Shuswap | Late; Fraser River | DU18 | 0.00% | 0.47% |
| | Little Lake | Late; Fraser River | DU18 | 0.47% | 0.00% |

| | | | | | |
|---|---|---|---|---|---|
| | Middle Lake | Summer; Fraser River | DU18 | 0.47% | 0.00% |
| | Middle Shuswap | Late; Fraser River | DU18 | 0.00% | 1.42% |
| | Mitchell River | Summer; Fraser River | DU16 | 0.47% | 0.47% |
| | Nadina River | Early Summer; Fraser River | DU8 | 0.00% | 0.47% |
| | Nimpkish River | Vancouver Island | | 0.00% | 1.90% |
| | Pinchi Creek | Summer; Fraser River | DU21 | 0.47% | 0.00% |
| | Pitt River | Early Summer; Fraser River | DU15 | 0.47% | 0.00% |
| | Quesnel Horsefly | Summer; Fraser River | DU16 | 0.00% | 1.90% |
| | Scotch Creek | Early Summer; Fraser River | DU18 | 0.47% | 0.47% |
| | Stellako River | Summer; Fraser River | DU7 | 0.00% | 0.47% |
| | Woss Lake | Vancouver Island | | 0.00% | 0.47% |
